# Task-related multivariate activation states during task-free rest

**DOI:** 10.1101/068221

**Authors:** Richard H. Chen, Takuya Ito, Kaustubh R. Kulkarni, Michael W. Cole

**Affiliations:** Center for Molecular and Behavioral Neuroscience, Rutgers University, Newark, NJ, USA; Behavioral and Neural Sciences Graduate Program, Rutgers University, Newark, NJ, USA

**Author notes:** **Corresponding author/Lead contact:** Richard H. Chen, 197 University Ave., Rutgers University, Newark, NJ 07102.

**Keywords:** dynamics, fMRI, graph theory, community detection, resting-state

## Abstract

Much of our lives are spent in unconstrained rest states, yet cognitive brain processes are primarily investigated using task-constrained states. It may be possible to utilize the insights gained from experimental control of task processes as reference points for investigating unconstrained rest. To facilitate comparison of rest and task functional MRI (fMRI) data we focused on activation amplitude patterns, commonly used for task but not rest analyses. During rest, we identified spontaneous changes in temporally extended whole-brain activation pattern states. This revealed a hierarchical organization of rest states. The top consisted of two competing states consistent with previously identified “task-positive” and “task-negative” activation patterns. These states were composed of more specific states that repeated over time and across individuals. Contrasting with the view that rest consists of only task-negative states, task-positive states occurred 40% of the time while individuals “rested,” suggesting task-focused activity occurs during rest. Further, analysis of task data revealed a similar hierarchical structure of brain states. Together these results suggest brain activation dynamics form a general hierarchy across task and rest, with a small number of dominant general states reflecting basic functional modes and a variety of specific states likely reflecting a rich variety of cognitive processes.

## Introduction

The brain is a distributed information-processing system with rich spatiotemporal dynamics underlying complex cognitive dynamics. A major goal of cognitive neuroscience is to create a mapping between these two forms of dynamics to better understand the neural basis of cognition. Recent insights in human neuroimaging research have improved this mapping by considering activity in more than one spatial location at a time. These approaches include multivariate pattern analysis (MVPA) of brain activity patterns corresponding to cognitive task events (Haxby et al. 2014; Haynes 2015), as well as functional connectivity (FC) analysis of brain network dynamics during rest and task (Fox and Raichle 2007; Smith et al. 2011; Hutchison et al. 2013). Building on these advances, here we use what we term “dynamic MVPA” (dMVPA) – MVPA applied to the temporal evolution of brain processes (Anderson et al. 2012; Betzel et al. 2012; Yuan et al. 2012; King and Dehaene 2014) – with functional MRI (fMRI) to more comprehensively characterize the repertoire of brain states across a variety of resting and task cognitive states.

In particular, we used a state space characterization of dynamics previously used to gain insight into other real-world complex systems (Junejo 2010; Furusawa and Kaneko 2012). This involves conceptualizing each whole-brain image in time as a single point in a high-dimensional feature space. Brain state changes are thus equivalent to movement through that state space. We primarily utilize distance between these points in state space, using spatial correlations (a standard distance metric) between time points to temporally cluster them into brain states extending through time. We perform further analysis using graph theoretical formulations. Relative to classical clustering approaches, graph theoretical community detection algorithms can better assign patterns near the edges of a cluster (Newman and Girvan 2004). This can provide a more comprehensive characterization of a state space’s large-scale organization.

Recent investigations into moment-to-moment changes in FC have been important for characterizing brain dynamics (Hutchison et al. 2013). Unlike these studies of FC dynamics, we identify activation pattern-based brain states, which are not limited in their temporal precision by the need for temporally extended windows. More importantly, identifying whole-brain activation pattern states (rather than FC/temporal covariance patterns (Smith et al. 2009, 2012)) facilitates the functional interpretability of those states. This is due to whole-brain fMRI activation patterns being more directly related to the vast fMRI task activation literature, which associates cognitive task manipulations with whole-brain spatial activation maps(Laird et al. 2009; Yarkoni et al. 2011). In contrast, previous studies relating resting-state FC to the fMRI activation literature focused on across-study covariance between brain regions (e.g., the similarity in activation level changes across all tasks) (Smith et al. 2009, 2012), rather than focusing on any specific activation patterns (e.g., the whole-brain activation pattern during the N-back task). These studies revealed resting-state networks that were similar to patterns of cross-study covariance, yet single brain activation states are known to activate multiple resting-state networks at once (Smith et al. 2012; Bzdok et al. 2016). Several recent studies have focused on resting-state activation patterns (Liu and Duyn 2013; Liu et al. 2013). However, these studies focused on relating these patterns to resting-state networks, rather than task activations. Here we compare resting-state activation patterns to specific task activation maps – potentially involving co-activation of multiple resting-state networks – to identify the likely functional relevance of spontaneous brain states during rest. We expected this approach to better characterize the organization of human brain states and potentially identify task-related states present during “rest” periods.

We propose the use of the term dMVPA given the similarity of this approach to existing MVPA approaches with fMRI and M/EEG. However, whereas standard MVPA uses supervised machine learning (Haynes 2015) to classify activation patterns during experimentally induced cognitive events, dMVPA can be applied to the moment-by-moment temporal evolution of brain states to characterize even spontaneous cognitive events. A well-established example of dMVPA is the identification of M/EEG microstates, as defined by spontaneous changes in the spatial configuration of the scalp electromagnetic field (Khanna et al. 2015). Critically, however, the identification of M/EEG microstates suffers from strong limitations on spatial certainty caused by the smearing of neuronal sources as the electromagnetic field is conducted to the scalp surface (Grech et al. 2008; Schoffelen and Gross 2009). This spatial smearing suggests identified M/EEG microstates are biased toward states that are spatially extreme (i.e., spatial distinct despite smearing) rather than both temporally and spatially distinct. Consistent with this, attempts to map M/EEG microstates to more spatially accurate fMRI data has yielded states with only limited similarity to well-known states from the vast fMRI task activation literature (Yuan et al. 2012). Thus, we use a complementary approach here with fMRI to identify more spatially precise brain states during rest. While others have recently begun investigating spontaneous activation patterns with fMRI (Liu et al. 2013; Chen et al. 2015a), these patterns have not been directly compared to task activation patterns. Importantly, as mentioned above, comparison to task activation patterns is a major advantage of this approach, since using fMRI allows comparison of spontaneous states to the vast fMRI task activation literature, which associates cognitive manipulations with whole-brain spatial activation maps (Laird et al. 2009; Yarkoni et al. 2011).

We primarily utilize rest data given the possibility that many brain states are visited in this unconstrained “task-free” context (Fox and Raichle 2007). This allowed us to obtain a broad sampling of possible brain states across many (N=97) individuals. We also supplemented this broad repertoire of spontaneous states with experimentally controlled states identified from a variety of tasks involving distinct cognitive functions. We hypothesized that activation pattern states form a hierarchical structure, with a small number of dominant high-level states reflecting basic modes of brain function and a wide variety of lower-level states reflecting the rich variety of cognitive states humans are capable of visiting.

In addition to this broad hypothesis, we sought to test the hypothesis that “task-positive” brain states occur regularly during rest periods. This contrasts with the highly replicated finding that a “task-negative” brain state with high default-mode network activity is present during rest (Greicius et al. 2003; Buckner and Vincent 2007; Fox and Raichle 2007). We expected that the task-negative state dominates rest (consistent with previous results), but that task-positive states also occur frequently. This expectation is based on the common self-reported experience of trying to solve the problems of everyday life during periods of “rest” (Smallwood and Schooler 2006). Such problem solving effort is associated with cognitive control demands, which involve task-positive activations in networks such as the frontoparietal control network and the cingulo-opercular network and deactivation of the default-mode network (Raichle et al. 2001; Cole and Schneider 2007; Anticevic et al. 2012). Additionally, everyday experience suggests some portion of “rest” consists of passive attending to external sensory and motor events, suggesting occasional activation of sensory/motor networks and attention networks (Fox et al. 2006). We expected that identifying the overall organization of spontaneous brain state dynamics would allow us to test the possibility of both task-positive and task-negative brain states occurring during rest periods, as well as allowing us to estimate the proportion of time devoted to each class of brain states.

## MATERIALS & METHODS

### Participants

Data were collected as part of the Washington University-Minnesota Consortium Human Connectome Project (Van Essen et al. 2013). The participants were recruited from the Washington University campus and surrounding area. All participants supplied informed consent. The data were from the “500 Subjects” public data release. We used data from the “100 Unrelated Subjects” set as we wanted a sample representative of the general population (excluding family relations).

We used resting-state fMRI and task-state fMRI data from 100 subjects, with 3 outlier subjects removed for a subset of analyses (see Results). The resting-state dataset consisted of four separate runs, each spanning 14.4 minutes in length. Analyses were performed separately for each rest run. The task data involved seven diverse tasks (Barch et al. 2013). These seven tasks were selected to tap into different cognitive processes as well as the different neural circuitry that supports those functions. The tasks were related to emotion perception, reward learning, language processing, motor responses, relational reasoning, social cognition, and working memory. See (Barch et al. 2013) for more details on the task fMRI data.

### MRI parameters

Whole-brain echo-planar scans were acquired with a 32 channel head coil on a modified 3T Siemens Skyra with TR = 720 ms, TE = 33.1 ms, flip angle = 52°, BW = 2290 Hz/Px, in-plane FOV = 208 × 180 mm, 72 slices, 2.0 mm isotropic voxels, with a multi-band acceleration factor of 8 (Uğurbil et al., 2013). Data were collected across two days. On each day 28 minutes of rest (eyes open with fixation) fMRI data was collected across two runs (56 minutes total), followed by 30 minutes of task fMRI data collection (60 minutes total). Each of the 7 tasks was completed over two consecutive fMRI runs. Details regarding the resting-state data collection for this dataset can be found elsewhere (Smith et al. 2013), as well as details about the tasks (Barch et al. 2013).

### fMRI preprocessing

We used a minimally preprocessed version of the data, which was the result of standard procedures including: spatial normalization to a standard template, motion correction, and intensity normalization. These steps have been described previously (Glasser et al. 2013). We performed analyses on the volume version of these minimally preprocessed data using AFNI (Cox 1996). We removed variables of no interest from the time series using linear regression, including: motion estimates, ventricle and white matter signals, as well as derivatives. Ventricle, white matter, gray matter, and anatomical structures were identified for each subject using FreeSurfer (Fischl et al. 2002, 2004). Note that whole-brain global signal was not removed due to controversy regarding this preprocessing step (Murphy et al. 2009). The linear trend was removed from the signal and the data were spatially smoothed (FWHM = 4 mm). Resting fMRI data is also typically temporally filtered to isolate the low frequency component of the time series. We did not apply a temporal filter to the data due to the possibility of relatively rapid brain state transitions.

Further data analysis was completed by sampling from a set of 264 regions in order to capture and explore large-scale regional and system-level questions. These 264 regions were independently identified in a previous study using functional criteria (Power et al. 2011). Using this approach reduces the chance of blurring signal from neighboring regions with different functional profiles (Wig et al. 2011). Specifically, the 264 regions were identified and classified using resting-state FC parcellation (Cohen et al. 2008) and a task-based neuroimaging metaanalysis (Power et al. 2011). The mean time series from all of the voxels within each of these 264 regions was then calculated and used in subsequent analyses. Subsequent data analyses were conducted with MATLAB 2014b (Mathworks) or Python.

### Brain state identification

We conceptualized time in terms of a weighted graph, with each time point as a graph node and edges as the similarities of whole-brain spatial activation patterns at those time points. Activation pattern similarity was calculated using Pearson correlation. Pattern similarity can be equivalently considered as (the inverse of) distance in state space, which has a long methodological history in mathematics and other fields (Cha 2007). Pearson correlation was used given that it is a well-established distance measure (Cha 2007), is invariant to scale changes (unlike, e.g., Euclidean distance), and can be conveniently conceptualized as (the square root of) linear variance explained. Note that Pearson correlation does not assume the underlying data are normally distributed when used as a similarity measure, given that its associated p-value is not calculated (Ahlgren et al. 2003). These correlation-based associations were summarized in a temporal similarity matrix, consisting of all pairwise similarities among time points.

Time points with strong edges between them were considered as instances of the same brain state. Clusters of similar time points were identified using the Infomap community detection algorithm (Rosvall et al. 2009; Rosvall and Bergstrom 2011; Bohlin et al. 2014). To determine the best clustering approach for brain state identification, we also tested two other distinct clustering approaches (k-means clustering and hierarchical clustering) on a single subject (to avoid potential overfitting of a specific method on the full dataset). We found that clustering solutions were quite similar to one another, with the Infomap solution performing slightly better based on modularity scores (see Supplementary for details). Infomap community detection algorithm was first applied to each subject’s initial resting-state fMRI run (14.4 minutes in duration, consisting of 1200 fMRI time points). Given the likely presence of noise, we thresholded the temporal similarity matrix at thresholds from 1% to 50% density (1% increments), prior to the application of Infomap. For each iteration of the density thresholding, we applied the Infomap algorithm to assign each time point to unique communities. Modularity was calculated to assess the quality of Infomap’s assignment of time points to communities (Blondel et al. 2008; Rubinov and Sporns 2011). A standard consensus approach (Lancichinetti and Fortunato 2012) was used to combined the assignments, weighted by the respective modularity scores, across all density thresholds (50 thresholds per subject) to obtain a final set of community assignments for an individual subject’s resting-state run. This produced on average 4 unique brain states (graph communities) per subject, with a total of 412 brain states across 98 subjects. Note that two of the 100 subjects were excluded because the Infomap algorithm returned more than three standard deviations above the average number of states per subject. We reduced the many time points contributing to each brain state to a single “prototype” vector for each brain state via averaging.

We next conducted a group-level analysis. This involved first computing an adjacency matrix based on the similarities among the brain state prototypes across all subjects. We then applied the Infomap algorithm to that matrix, this time with no thresholding, to produce a group-level brain state solution. Later, we tested for the presence of a hierarchy of brain states, with the no-threshold solution being the top of the hierarchy. We produced lower levels of the hierarchy by re-running the algorithm (independently of the previous level) using a series of thresholds. We began with 100% density (no threshold), going down by 10% increments until the 10% density level. In contrast to the subject-level analysis, no consensus approach was applied across the density thresholds. Each density threshold was analyzed independently, since reducing the density eliminates weaker connections and allowed for better isolation and separation of the communities (Power et al. 2011).

### Task relevance – correspondence between brain states and behavioral/mental states

We sought to identify correspondences between the identified brain states and known behavioral/mental states based on task manipulations. This allowed assessment of the likely functional relevance of the identified brain states. Task-rest correspondence was assessed using pairwise Pearson’s correlations between each group-level brain state and every activation pattern in each time point across all subjects for both resting fMRI and task fMRI. This returned a time series of correlation scores between each subject and the group-level brain states. The time series of all subjects were then visualized and assessed for temporal dynamics of brain state transitions.

### Assessing the amount of time spent in each brain state

For resting-state data, each individual’s Infomap clustering solution was re-mapped into a two-state solution based on the group-level clustering. The durations in each state were then calculated based on this two-state solution for each individual subject. For task-state data, each time point was classified as either State A or State B using a support vector machine (SVM) classifier trained on resting-state data. Specifically, the SVM classifier was trained on rest runs 2-4 and validated by testing on rest run 1 prior to classification of task state data (for each task separately). Note that validation with rest run 1 data was based on above-chance classification accuracy, using the InfoMap clustering labels as the ground truth. The two-state classification results were used to determine the amount spent in each state for each of the seven tasks.

### Neurosynth state decoding

Brain states were decoded using the Neurosynth decoder tool (Yarkoni et al. 2011). Neurosynth is a meta-analytical tool that contains a database with brain activation patterns and peak signal coordinates paired with the associated cognitive terms from the fMRI scientific literature. The decoder function takes in our voxelwise representation of brain states, cross-references with the database, and returns a list of cognitive terms each paired with a correlation score indicating how well each brain state is associated with each cognitive term. The decoder returned a list of approximately 3400 cognitive terms and their correlation values with the tested brain state. Of the top 50 highest correlated terms, all anatomical terms, redundant terms, nonsensical terms (i.e., “cortexmpfc”, “networkdmn”), and methodological terms were excluded from the list. The remaining terms were visualized as word clouds and the relative font sizes of each term were determined by the correlation scores.

We sought to validate the results returned by Neurosynth by running a series of permutation tests for each brain state. Specifically, within each subject, we permuted the order of the brain state temporal windows (consecutive time points of the same state) as returned by the Infomap clustering process. This approach allowed us to preserve many of the temporal properties of brain states within each subject, while still permuting the data being averaged for each unique brain state label. The newly “permuted” brain state prototypes were averaged across subjects according to the group-level clustering solution to produce a “permuted” State A and State B. The “permuted” State A and State B were run through the Neurosynth decoder to obtain correlation scores for each of the 3400 terms. A total of 1000 permutations cycles were performed for each brain state, returning a total of 340,000 correlation scores. This allowed us to create a null distribution of r-values to test the statistical significance of the observed r-value for each cognitive term. Note that these permutation tests controlled for multiple comparisons, since all 3400 comparisons were computed during each permutation (Nichols and Holmes, 2002). The same analysis was also performed for each of the 12 states at the 20% density tier of the hierarchy (see Supplementary for methods, detailed results, and figure).

## RESULTS

### Identifying multivariate brain activation states

We sought to characterize the repertoire of spontaneous human brain activation patterns using a data-driven method of temporally clustering resting-state fMRI activity. We hypothesized that whole-brain fMRI activation patterns would consist of discrete “states” – configuration patterns that extend (and repeat) through time. To test this possibility we clustered whole-brain resting-state fMRI activations in time. A standard set of functionally defined regions (Power et al. 2011) were used for computational tractability, as well as for the previously identified functional system assignments (to potentially increase interpretability of observed states) (Figure 1D). An alternative parcellation set (Gordon et al. 2014) was also used to test the replicability of the observed brain state results (see Supplementary for details). Spatial correlations were used as a similarity/distance measure across individual time points, with brain states defined as temporal clusters of similar activation patterns (Figure 1A).

**Figure 1.**
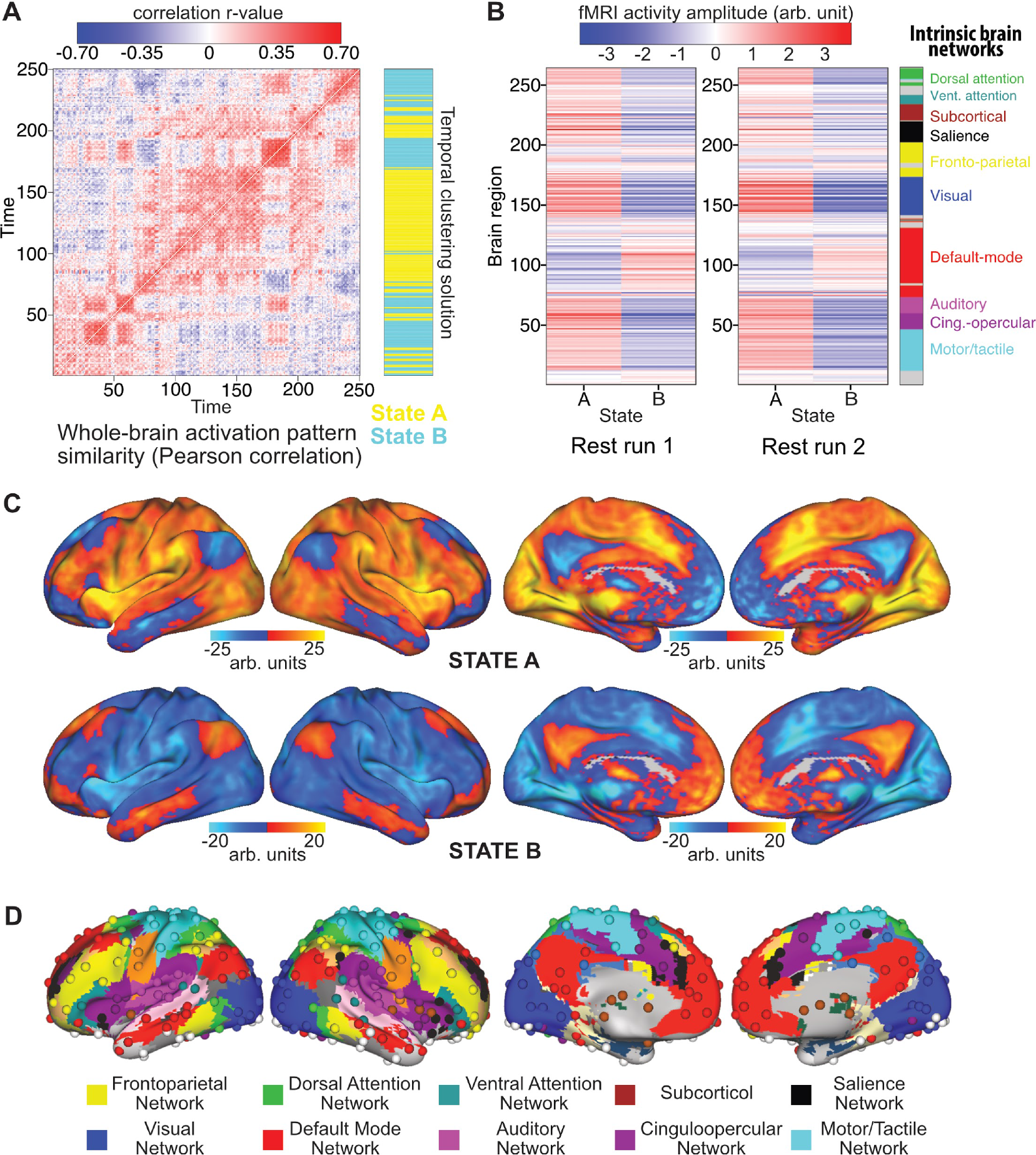
The repertoire of brain states based on resting-state fMRI. Each individual’s prototypical brain states are correlated and clustered using the same Infomap algorithm. **A**. An example of individual time point similarity based on spatial correlations. 250 time points from a single subject (HCP subject 100408, time points 100:350) are shown for illustration. The across-subject clustering result (mapped back to this subject’s data) is shown in blue and yellow. **B.** Brain state prototypes for state A and state B, averaged across all activation vectors across all subjects. The two-state results replicated across all 4 of the rest runs (results from 2 runs are shown in **B**). **C**. Voxelwise representations of state A and state B. Note that state B involves activation of the default-mode network as well as the tan/salmon-colored portion of the frontoparietal network in panel D. **D**. Functionally defined set of 264 regions and the associated functional network assignments.

Brain states were identified first at the individual subject level. Of the 100 subjects, two subjects were excluded from subsequent analyses (unless noted otherwise) because the clustering algorithm returned more than three standard deviations above the average number of states per subject. On average, four unique brain states were identified for each subject for a total of 412 unique brain state prototypes across the remaining 98 subjects. These 412 brain states were re-clustered in a group level analysis. The same processing steps performed at the individual subject level were applied at the group level for determining group-level brain states (see Methods for details).

At the group level two brain states were identified, which we labeled “State A” and “State B” (Figure 1B & 1C). The states were summarized by averaging all prototypes with the same clustering label and visualized in 264 ROI space (Figure 1B) and in voxel space (Figure 1C). An example subject’s temporal similarity matrix along with the group-level clustering solution is illustrated in Figure 1A. Periods of highly similar activation patterns can be observed by the blocked structure on the diagonal of the temporal similarity matrix, which are grouped as the same state in the clustering solution. We replicated the current findings using the same approach for the remaining three rest runs and found similar results (second run shown in Figure 1B). Specifically, State A identified in any given rest run was highly correlated on average with State A identified in other rest runs: rank correlation rho=0.92 (p<0.0001 for all pairwise comparisons). This was also the case for State B being correlated with State B identified in other rest runs: average rank correlation rho=0.91 (p<0.0001 for all pairwise comparisons). Note that we used Spearman’s rank correlation (rho) to calculate p-values for data that were not approximately normally distributed. Additionally, State A and State B were highly anti-correlated at rho=-0.97 on average (p<0.0001 for all pairwise comparisons). Note that the clustering approach used to identify distinct activation states prioritizes identification of maximally distinct (e.g., anti-correlated) states. To confirm the statistical significance of the observed anti-correlation despite this prioritization of maximally distinct states, we performed a permutation test with 10000 iterations, finding that the observed result was p<0.0001 (see Supplementary section for methods and detailed results).

Functionally, State A is highly similar to observed “task-positive” activation patterns reported in the literature (Corbetta and Shulman 2002; Fox et al. 2005; Power et al. 2011). This activation pattern includes active regions in many task-related functional networks, such as the dorsal and ventral attention networks, salience network, and sensory-motor networks (visual, auditory, somatomotor). Conversely, State B may be a “task-negative” state, given the strong activation of the default mode network (DMN) (Fox et al. 2005; Uddin et al. 2009) (along with a portion of the fronto-parietal cognitive control network (FPN)). Note, however, that we used only “task-negative” data – fMRI data collected during rest – to define both of these states.

### Spatial activation patterns vs. FC

Using resting-state data, we identified a “task-positive” state A and a “task-negative” state B. These labels were based on the functions assigned to these activation patterns by the task activation fMRI literature (Duncan and Owen 2000; Raichle et al. 2001; Smith et al. 2009; Laird et al. 2013). As mentioned above, several aspects of these state A and state B patterns (Figure 1B) are consistent with networks identified in the resting-state FC literature (Fox et al. 2005; Dosenbach et al. 2008; Power et al. 2011). However, the approach used here was quite distinct from FC analyses, since it is based on spatial activation covariance, which is ignored by FC approaches in favor of temporal covariance. Therefore, in order to test whether the current spatial activation pattern approach reveals additional insights independent of traditional FC analyses, we analyzed resting-state FC patterns across time periods identified as either State A or State B (Fig 3). Comparing between the FC patterns calculated by averaging across all subjects (Figure 2), we found that the overall resting-state FC pattern (calculated across the entire rest run) correlated with State A FC at r=0.948 and State B FC at r=0.951. Additionally, to test whether results were consistent across subjects, FC comparisons were also performed for each subject separately. We then performed random-effects statistical analyses to test for across-subject consistency. The overall resting-state FC pattern correlated with State A FC on average at r=0.55 (t(96)=42.4, p<0.0001) and State B FC on average at r=0.54 (t(96) = 49.1, p<0.0001). Note that we calculated the overall resting-state FC pattern based on a separate run (e.g., run 2 when comparing to run 1’s State A FC) to remove circularity from the analysis (Kriegeskorte et al. 2009). All non-circular combinations (between the first and second resting-state run) were computed and the reported results are based on averages across these comparisons.

**Figure 2.**
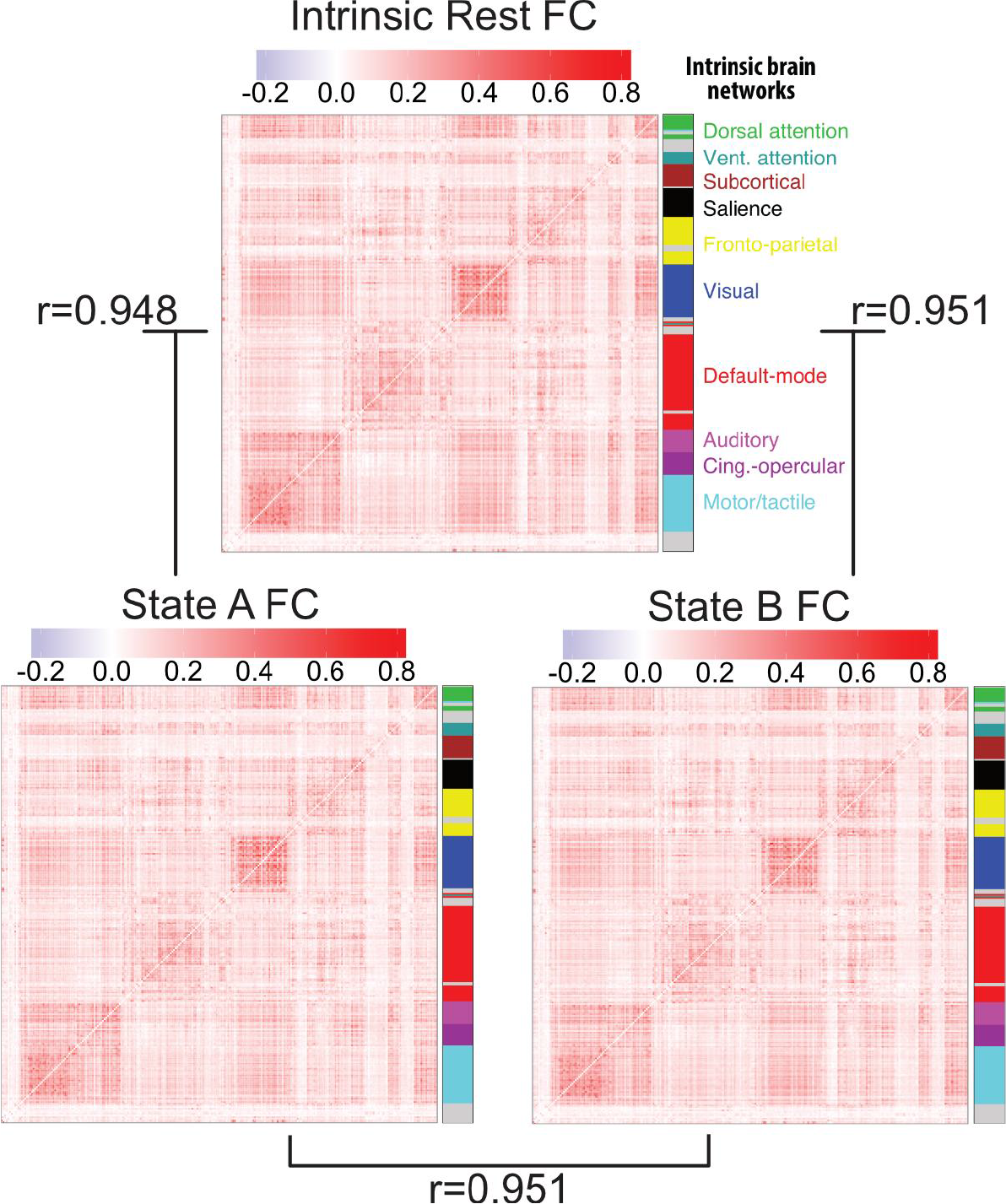
Independence of activation state dynamics from FC state dynamics. The FC matrices for resting-state (over the entire rest run), State A (just the time points labeled state A), and State B (just the time points labeled state B) are pairwise correlated with each other for each subject. Averaged FC matrices and correlation values across subjects are shown here. These results suggest that the whole-brain multivariate activation states investigated here are independent of FC dynamics.

**Figure 3.**
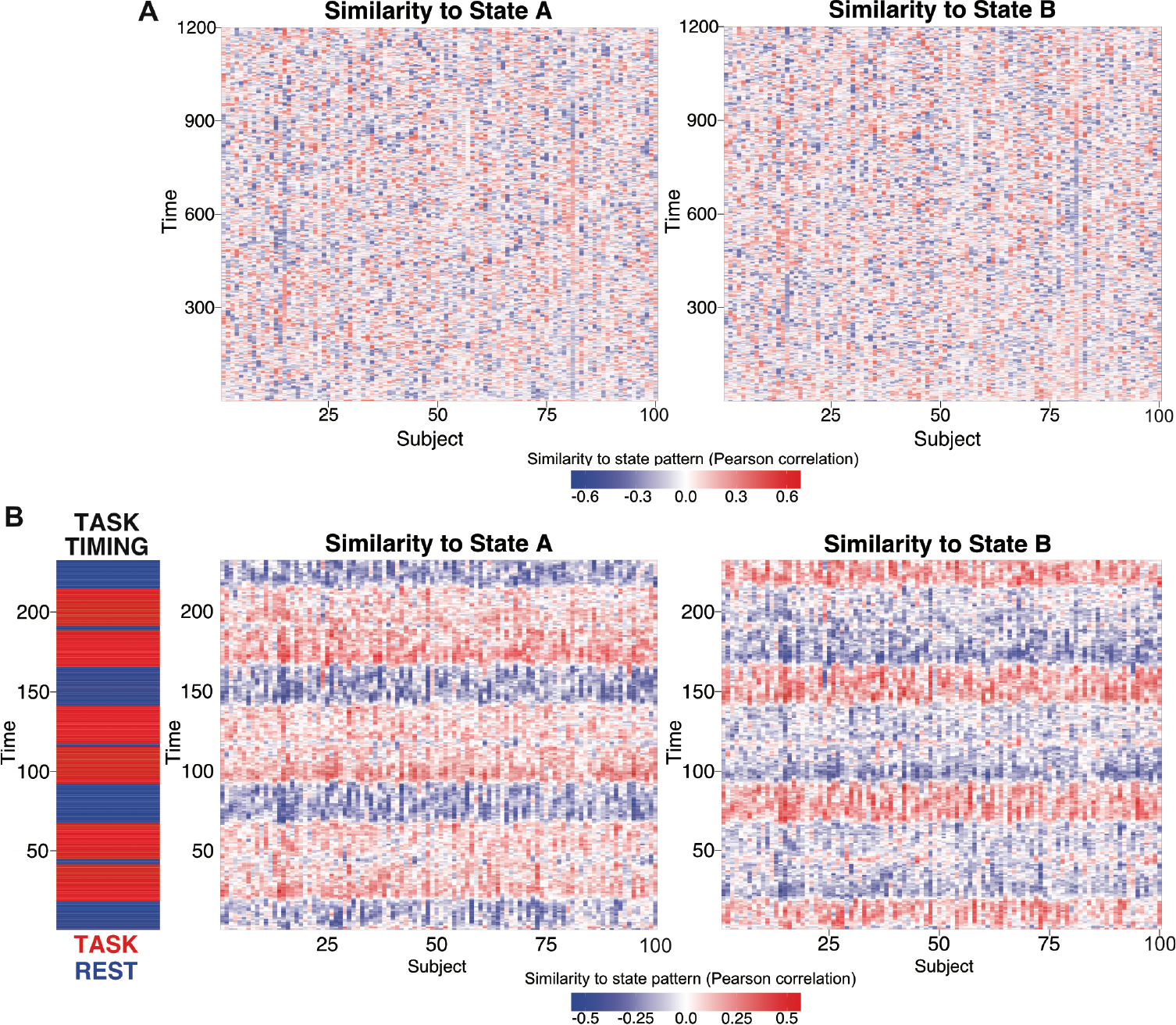
Functional relevance of the two main states.

The two brain states observed at the group level (see Figure 1B) are correlated with each subjects’ rest and task data (separately for each time point). No discernable temporal organization of state A and state B is observed in resting-state data across all 100 subjects (**A**). The two subjects removed during the state identification analysis were included here for completeness. Evidence of state A during rest may suggest subjects are performing covert tasks during rest. Clear structure is seen when correlated with activity during the reasoning task (**B**). All subjects consistently enter state A when entering task blocks (in red), and enter state B when entering inter-task rest periods (in blue).

The strong correlation between the individual states’ FC and overall rest FC suggests that resting-state FC architecture remains consistent across distinct activation brain states. Further supporting this conclusion, across-subject averaged State A FC correlated with across-subject averaged State B FC at r=0.951. This was also the case when computed based on averaging within-subject comparisons: r=0.49 (t(96) = 48.6, p<0.0001), suggesting across-subject consistency. Note that this was calculated by comparing across rest runs, as with the overall resting-state FC pattern comparisons above. The resulting average similarity is very high relative to chance, but also in contrast to the strong anti-correlation (rho=-0.97) between State A and State B activation patterns. This suggests the possibility that whole-brain activation pattern dynamics are largely independent of FC dynamics. This supports our choice to analyze activation patterns independently of FC, as well as identifying brain states based on activation pattern similarities rather than FC dynamics.

### Functional relevance of the two states

Identifying a task-positive State A when using “task-negative” resting-state data suggests that subjects may have been performing covert tasks during “rest” periods. To test this hypothesis, we compared both State A and State B with rest and task data to better understand the functional significance of the two brain states. Specifically, we explored the temporal characteristics of the two states and how they related to ongoing task demands by correlating the two states’ prototypical activation patterns (Figure 1B) with every individual time point’s whole-brain activation pattern. This was done for all subjects individually in both resting-state and in each of the seven tasks (Figure 3B shows correlation results for the reasoning task). No across-subject temporal patterns were observed in the correlations with resting-state data (Figure 3A), as expected. On average, subjects were in State A only 39% of the time, and in State B 61% of the time. In contrast, for each of the seven tasks (Table 1), we found that every subject was in State A most of the time during task blocks and in State B during the inter-task rest periods (Figure 2B), with the exception of the language task (see Discussion). On average (including the language task), subjects were in State A 54% (State B 46%) of the time during task blocks and in State A 41% (State B 59%) of the time during inter-task rest periods. Additionally, subjects tended to stay in State A longer for tasks that were likely more challenging. For instance, subjects were in State A 59% (State B 41%) for the reasoning task, but only in State A 49% (State B 51%) for the language task. This suggests that State A is not only relevant to rest data (the data used to derive it), but also likely to be the same state required for performing various active task demands.

**Table 1.** Percent time in each state for each task. Percent of time in each state during task blocks and inter-rest rest periods for each task.

|  | Task Blocks |  | Inter-task Rest Blocks |  |
| --- | --- | --- | --- | --- |
| Task Type | State A | State B | State A | State B |
| Reasoning | 59% | 41% | 36% | 64% |
| Emotion | 52% | 48% | 40% | 60% |
| Working Memory | 53% | 47% | 39% | 61% |
| Gambling | 57% | 43% | 38% | 62% |
| Language | 49% | 51% | 50% | 50% |
| Social | 55% | 45% | 41% | 59% |
| Motor | 53% | 47% | 40% | 60% |
| *Average* | *54%* | *46%* | *41%* | *59%* |

### Hierarchical organization of brain states

The previous analyses suggest that State A and State B may be general and multifunctional states. Further, the vast fMRI literature indicates that each active task (cognitive state) has a unique activation pattern, despite an overarching “task-positive” activation pattern for externally oriented and cognitive control tasks (Corbetta and Shulman 2002; Fox et al. 2005; Power et al. 2011). We therefore hypothesized brain states have a hierarchical organization, with state A and state B at the top level and more specific task states at lower levels (Figure 4). The two top states were broken down into more states by reducing the density of the adjacency matrix before applying Infomap clustering, from 100% to 20% density at decreasing increments of 20%. Reducing the density of the adjacency matrix removes weaker graph edges (lower correlations) between the group state prototypes. The task-positive state A divides into two states early in the hierarchy (60% density), shown by the red links between the levels of the hierarchy (Figure 4). The task negative state B splits into multiple states at the lowest level of the hierarchy (20% density), shown with blue links between the levels of the hierarchy.

**Figure 4.**
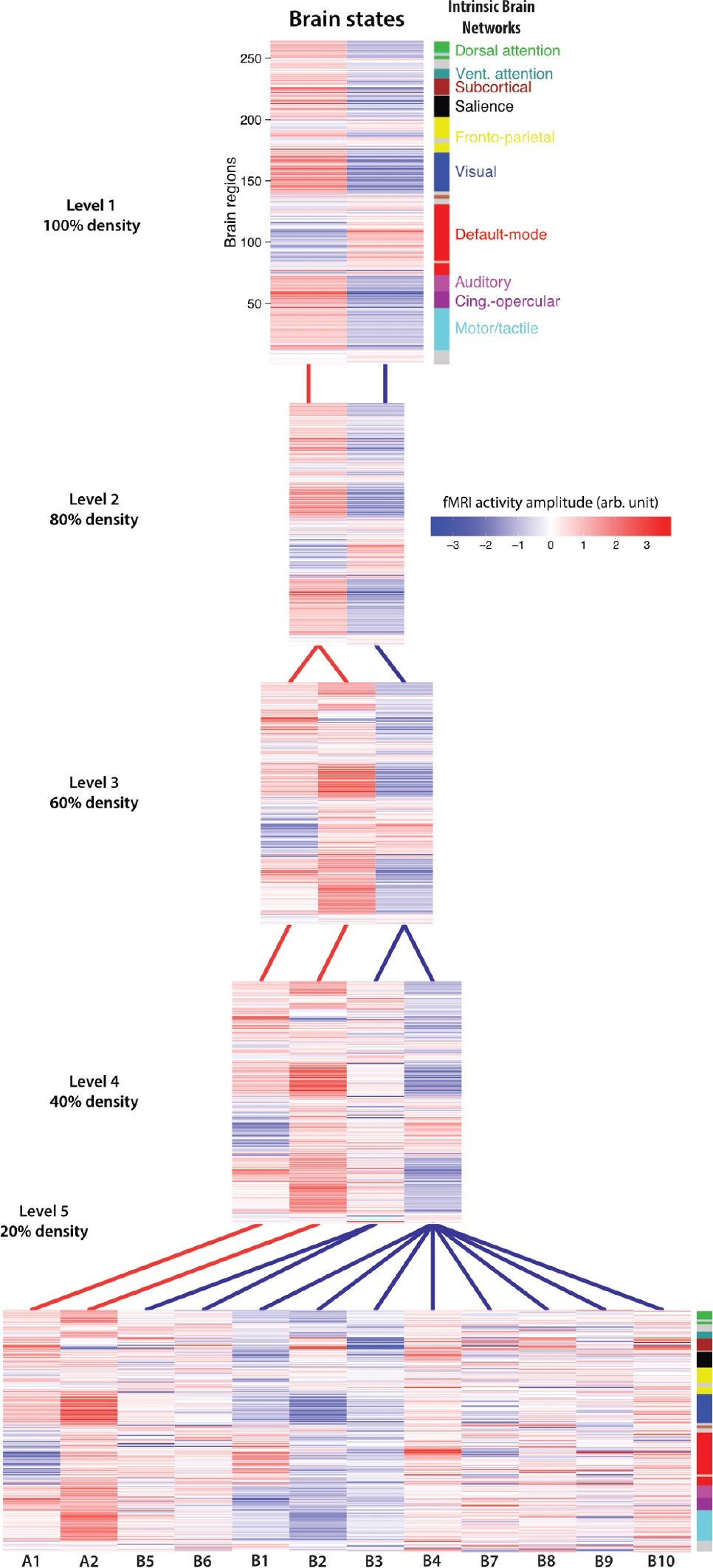
Relationship between levels of the brain state hierarchy. The top level of the hierarchy (i.e., with no thresholding) is depicted at top, with additional states lower in the hierarchy with more stringent thresholding. Each level is defined by removing graph edges between the group prototype states (e.g., 80% density means removing the weakest 20% of edges) prior to running the community detection algorithm. The lines depict which state in the higher level that the lower-level state is most associated with. Specifically, a line indicates which higher-level state is composed of the largest percentage of the same group-level prototypes as the lower level state. Red links are associated with State A (task-positive) at the top level while blue links are associated with State B (task-negative) at the top level. The state labels below each of the states at 20% density (level 5) corresponds with the state labels in Figure 7.

To better understand the relationships between the individual prototypes that contribute to the state hierarchy, we visualized all 412 individual state prototypes in a force-directed graph (Figure 5). This helped characterize the state space underlying the states in the hierarchy. Each node in the graph represents an individual’s brain state prototype. Edges connecting the nodes were weighted based on Pearson correlation scores between each pair of nodes, with stronger weights pulling nodes closer together. The correlation strengths are depicted on the graph with solid lines (p<0.01) and dashed lines (p<0.05). Negative correlations were excluded as possible edges on the graph. The nodes are colored based on the Infomap clustering assignments for the 100% density level (Figure 5A) and the 20% density level (Figure 5B). At the top level of the hierarchy, the centers of the two states consist of tightly clustered and strongly connected prototypes. However, at the 20% density level, the nodes located further away from the centers of State A and State B break off to form several clusters of prototypes, each constituting a unique brain state. Five states with the largest clusters are emphasized in Figure 5B. Two of these clusters (A1, A2) are from state A and three of the clusters (B1, B2, B3) are from state B.

**Figure 5.**
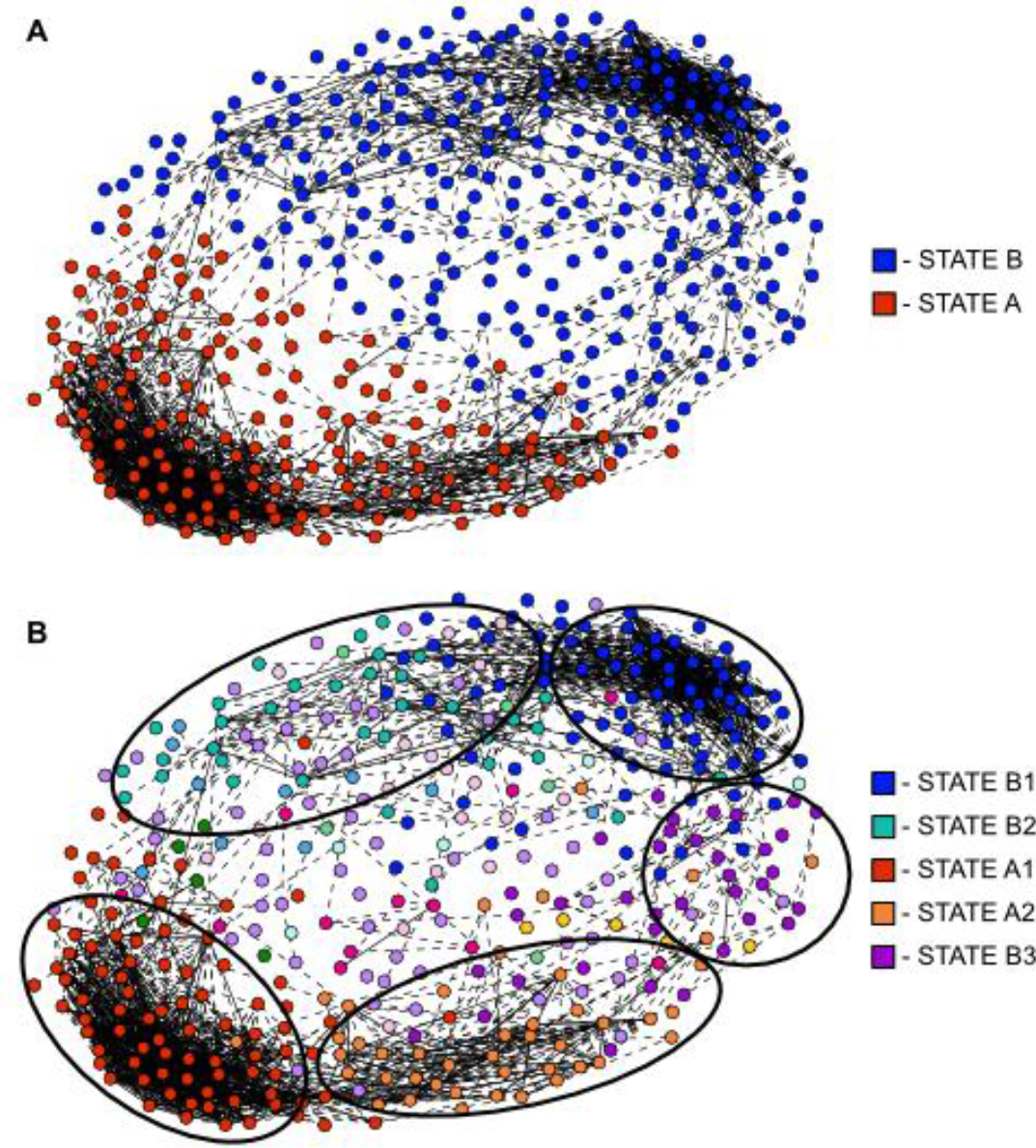
Force directed graph visualization of brain states. All 412 group-level prototype brain states are depicted as nodes in a force-directed graph. More spatially similar states are closer together, while less similar states are forced apart. Solid lines indicate a significant spatial correlation between states at p<0.01, while a dashed line indicates a significant correlation at p<0.05. Panel **A** indicates the top-level solution (100% density), with red being the task-positive State A and blue being the task-negative State B. Panel **B** shows the lowest level of the hierarchy (20% density), with each color indicating a separate graph community. Large communities are highlighted with circles. Note that states appear to break off from the primary State A and State B clusters when they are mixtures of those two states (i.e., they were between the large State A and State B clusters).

### Hierarchical organization in task fMRI

We next analyzed task-state fMRI brain states to test the hypothesis that task activations have a similar hierarchical structure as identified using resting-state fMRI data. Importantly, we could use a more supervised approach here, since the experimental cognitive manipulations indicate (roughly) which mental state each subject is in. This involved assuming the experimental cognitive state corresponded to the brain activation state, then working backward to see if the activations were hierarchical in nature.

We first calculated the mean brain state prototypes for each of the seven tasks, shown in Figure 6A. This was done using a standard fMRI general linear model, with a canonical hemodynamic response function (given that all 7 tasks used block designs) and inter-block rest periods as baseline. The extremely high similarity across the seven distinct cognitive states observed (r=0.9984 on average) suggests that the highly similar brain states identified with resting-state fMRI may map to distinct cognitive states as well.

**Figure 6.**
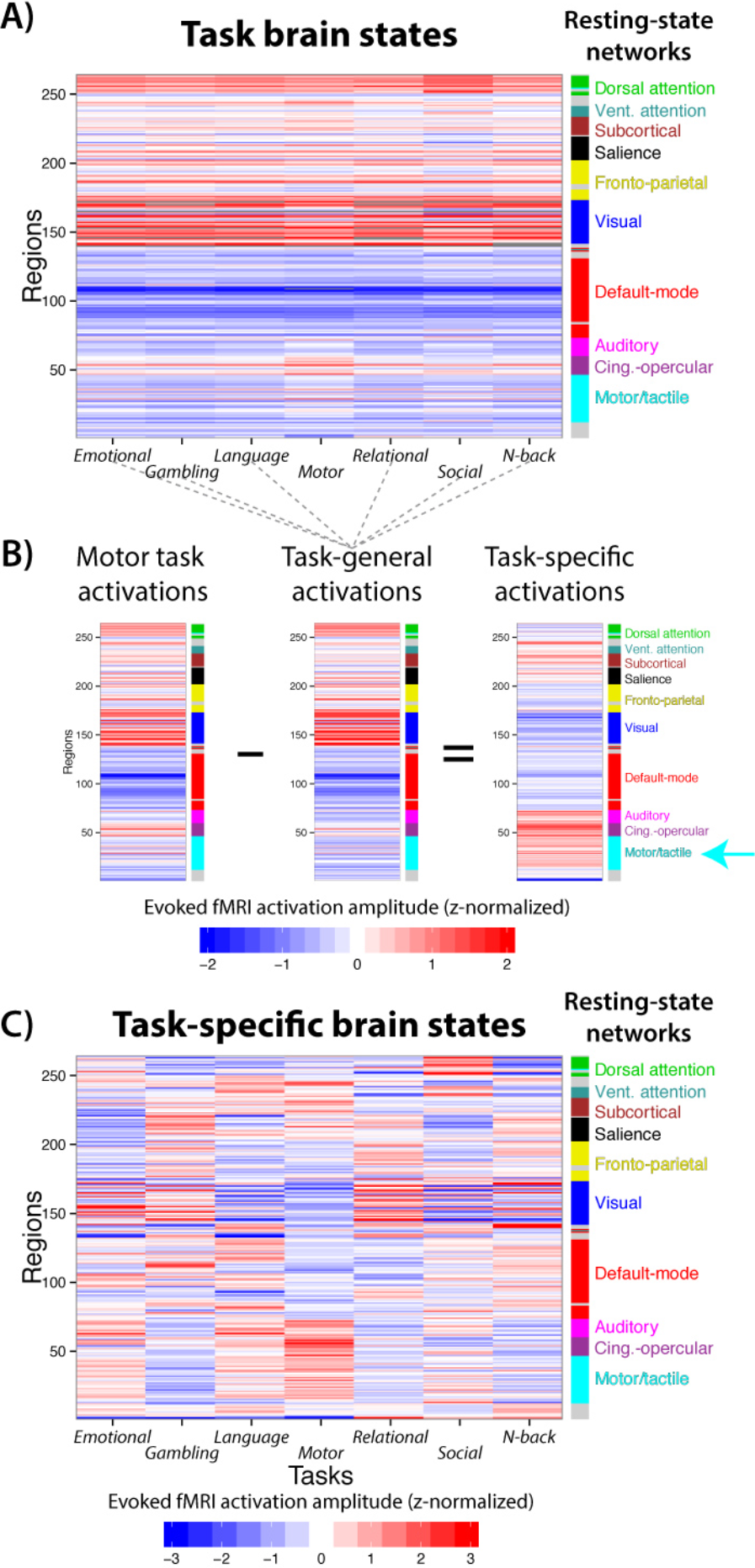
Hierarchical organization of task activation brain states. **A**) The mean whole-brain activation patterns for each state are shown. These brain states were highly similar to each other on average (r=0.9984). The similarity of the brain states underlying these quite distinct mental states suggests the similarities observed across the resting-state brain states likely correspond to distinct mental states as well. **B**) We hypothesized that task activation states are also organized hierarchically. We used PCA to derive a task-general activation pattern – a higher-order brain state – which was correlated with State A (r=0.56, p<0.0001) but anti-correlated with State B (r=-0.54, p<0.0001). A lower-order task-specific brain state was revealed when the higher-order task-general brain state was subtracted from each task’s activation pattern. The motor task is shown for illustration. Motor/tactile network activations are not readily apparent in the original activation pattern, but are revealed when the task-general pattern is subtracted. Note that such task-specific patterns are typically observed using specific contrasts (e.g., left vs. right hand motor responses), whereas we use a task-general contrast here. **C**) Task-specific activation patterns for all 7 tasks are shown (average across-task r=-0.01). These results suggest important functional distinctions relevant to cognition and behavior are hierarchically embedded within the observed rest states despite dominance of State A and State B.

Next, we used principal component analysis (PCA; the first principal component) to isolate a dominant task-general activation state (Figure 6B middle), which was quite similar to the State A pattern identified using resting-state data (r=0.56, p<0.0001) but anti-correlated with State B (r=-0.54, p<0.0001). The task-general activation state was then removed from each of the seven mean task states using linear regression to obtain task-specific activation states (Figure 7B right). Each of the task-specific activation states included active regions associated with each cognitive task. For example (Figure 6B), after subtracting the task-general activation state from the motor task state, the remaining task-specific activation state included predominantly motor region activity and some attention region activity. Similar results were observed for the remaining six tasks as well (Figure 6C). The presence of a strongly dominant task-general state across diverse task demands, along with subordinate task-specific activation states across those tasks, indicates a hierarchical organization of the state space that is similar to what was observed with resting-state fMRI.

**Figure 7.**
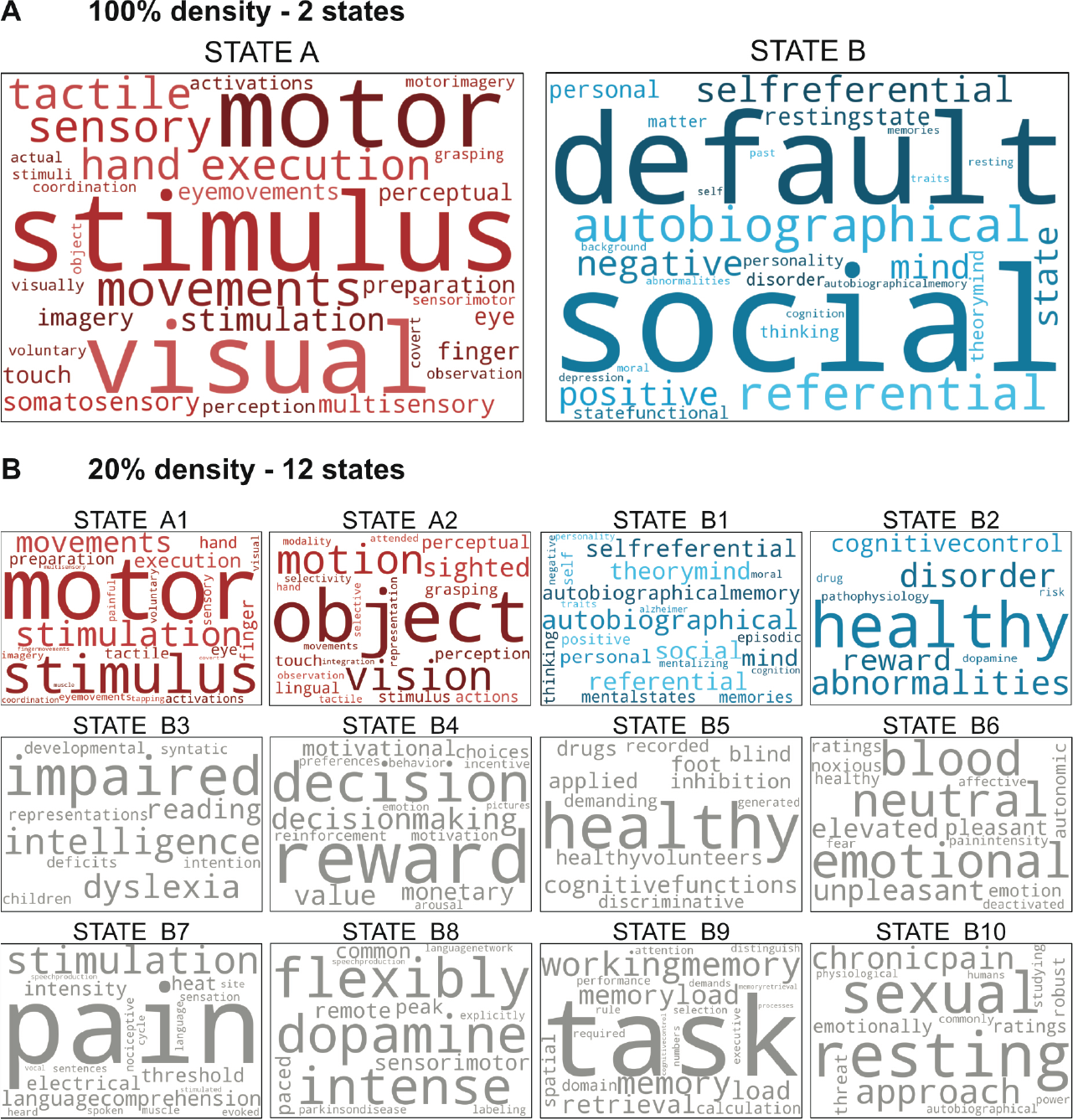
Decoding the states using Neurosynth. Each state at the 100% (A) and 20% (B) levels of the hierarchy illustrated in Figure 6 were decoded using Neurosynth. Cognitive terms with the strongest associations with each state are depicted as word clouds (top 50 terms with non-functional terms removed; see Methods for details on term selection). Terms with higher correlation values with the indicated brain state activation pattern are represented by larger font sizes (with size normalized within each brain state’s word cloud). States B3-B10, colored in grey, did not survive the permutation testing (see Supplementary for details).

### Decoding brain states using Neurosynth

We next sought to take advantage of the vast fMRI task activation literature to decode the identified resting-state brain states, identifying likely cognitive processes occurring during rest. This also allows us to test the hypothesis that State A and State B are likely to be general states governing basic modes of cognition, while the lower level states (12 states at 20% density) are likely more functionally specific. To examine the specific cognitive functions of each state at the 100% and 20% density levels, each of the 14 states (state A, state B, and the 12 states at 20%) were decoded using the Neurosynth decoder function and visualized as word clouds. The word clouds for state A and state B (Figure 7A) were consistent with our previous interpretation of a “task-positive” state and a “task-negative” state, respectively. Additionally, the cognitive terms for each of the lower level states (Figure 7B) are quite distinct from one another, despite certain terms repeating across states (“healthy” for state B2, B5). State A branched into two lower-level states that shared similar cognitive terms. In contrast, State B is subdivided into ten lower-level states that include more diverse cognitive terms spanning multiple functional domains not seen in State B word cloud. One possible explanation for the variability in cognitive terms observed could be that State B is also an internally-focused state for passive homeostatic functions and covert task executions, which could contribute to a wider spread of equally significant cognitive terms in the lower-level State B word clouds (see Discussion).

We validated the Neurosynth findings by running a permutation test for each of the brain states (see Methods for details). We found that all of the cognitive terms of interest depicted in State A and State B word clouds were significantly different when compared against the permutation tests’ null distributions (p<0.001) for all cognitive terms (see Supplementary Figure 2). Additionally, all cognitive terms for State A1, A2, B1, and B2 were significantly different (p<0.05) when compared to the null distributions. Note that the permutation tests controlled for multiple comparisons, since all 3400 term comparisons were included in each permutation (see Methods). The remaining States B3-B10 (Figure 7B, in grey) did not survive the permutation tests. Together, these results support the accuracy of Neurosynth decoding of spontaneous activation states and suggest that the identified brain states correspond with distinct mental/cognitive states.

## DISCUSSION

By characterizing moment-to-moment activation patterns in a high-dimensional state space, we identified a hierarchical organization of functional brain states. Two domain-general states (State A and State B) occupy the highest tier of that hierarchy. These two states can be further subdivided into functionally specific “sub-states”. As expected, subjects spent a majority of time in State B, which matches the activation pattern commonly observed to be more active during rest than during task. Despite using resting-state data for brain state identification, we found that subjects spent a significant portion of time (39%) in State A, which matches the activation pattern commonly observed to be more active during a wide variety of tasks than during rest. The functional relevance of these activation patterns was characterized using a variety of distinct task states as well as pattern decoding based on meta-analysis of thousands of fMRI studies. Together these results suggest that whole-brain activation state configurations correspond between rest and a variety of tasks.

### Hierarchical Organization of Brain States

We identified a pair of competing anti-correlated states based on spontaneous activation patterns. State A appears to be a “task-positive” state, with high activation amplitudes in common task-active areas such as sensorimotor and cognitive control networks (Corbetta and Shulman 2002; Cole and Schneider 2007; Dosenbach et al. 2007; Raichle 2010). State B appears to be a “task-negative” state, with high activation patterns mainly in the default-mode network (Raichle et al. 2001; Buckner et al. 2008). Breaking individual brain states into more clusters, we observed 12 distinct brain states that are more functionally specific than State A and B, as indicated by associated terms from the cognitive neuroscience literature. Qualitatively, the activation pattern for each brain state includes highly activated and/or de-activated brain regions that are affiliated with various functional network definitions (Figure 4 bottom), further supporting the specificity of each brain state. However, detecting the same states across unconstrained rest and task at lower tiers of the hierarchy would be unlikely, given that any two subjects are unlikely to enter the exact same mental state spontaneously. Additionally, reverse inferences are more likely to be problematic for lower-level (more specific) states, since reverse inferences are often used inaccurately when making overly broad generalizations about specific activations (Poldrack and Wager 2006). Therefore, we focused predominantly on State A and State B, which were both present for 97 of the 98 subjects.

### Relation to Resting-state FC

In this study, we emphasized the use of whole-brain activation patterns over alternative FC-based approaches (Hutchison et al. 2013; Allen et al. 2014) for several reasons. Mainly, by focusing on activation amplitudes, we were able to directly compare the identified patterns with the large fMRI activation amplitude literature (Yarkoni et al. 2011; Laird et al. 2013). Also, characterizing activations allowed us to directly test whether subjects were in active task (externally oriented) brain states during “rest” periods, which the results support (Figure 7). Importantly, we found that FC architecture remains relatively unchanged across State A and State B despite significant differences in the underlying activation patterns (Figure 2). This suggests the activation states identified here are only weakly (if at all) related to FC states (Hutchison et al. 2013; Allen et al. 2014). However, the observed activation states appear to be related to static resting-state FC networks (based on estimating FC across entire resting-state runs). For instance, the State B activation pattern is highly similar to the DMN resting-state network, though a portion of the FPN is also present. Several studies (Liu et al. 2013; Chen et al. 2015b) have investigated the relationship between static resting-state networks and co-activation patterns more directly. These studies showed that co-activation patterns can resemble common resting-state networks identified using traditional FC-based approaches, but some differences in the co-activation patterns remained undetected in FC-based analyses. Together, these observations support the use of activation patterns to investigate brain state organization.

Despite evidence that the present results are independent of FC dynamics, these results appear to be related to observed anti-correlation between DMN and task-positive network time series according to (static) resting-state FC estimates (Fox et al. 2005). That result has been recently questioned, however, given that it is dependent on the global signal regression preprocessing step (Murphy et al. 2009). Importantly, we did not use global signal regression here, such that DMN time series were not anti-correlated with task-positive network time series (see Figure 2). This suggests the observed spatial (not temporal, which is used with FC) anti-correlation between DMN-dominated and task-positive-dominated states are at least somewhat independent of FC-based results. This may have been possible because spatial anti-correlation does not imply temporal anti-correlation (and vice versa). For example, two networks can be activated above baseline (high spatial correlation across time points) while being temporally anti-correlated at higher frequencies. It remains unclear exactly how spatial activation dynamics relate to time series dynamics, however, such that it will be important for future studies to fully characterize the relationship between spatial and temporal brain activity correlations in the future.

### Brain Activation States Common Across Task and Rest

By using a data-driven approach across dozens of subjects we were able to obtain a comprehensive characterization (at the spatiotemporal scale of multiband fMRI) of brain states across rest and task. The results confirmed our hypothesis that “task-positive” brain states occur regularly during rest periods. We specifically found that subjects spent the majority of rest in State B (61% of the time) but that a significant portion of rest was also spent in State A (39% of the time.) This ratio is reversed (as expected) during the seven tasks: subjects were in State A more often (54% of the time on average) during task periods. The only exception was the language task, where subjects were in State A only 49% of the time during task periods. One potential explanation is that the language task includes two trial types: a math task and a story task. It is possible that during the story task, subjects enter a self-reflective cognitive state consistent with State B, either due to the introspective nature of stories or due to a lack of active task demands, which may bias the percentages in favor of State B over State A during task periods.

A popular account of resting periods is that mind wandering is the primary mental phenomenon occurring during those times. Past studies have linked increased activity in the DMN, present in the State B activation pattern, with mind wandering (Mason et al. 2007; Mittner et al. 2016). However, it is likely that increased DMN activity is not the only neural mechanism underlying mind wandering. The significant presence of State A suggests that the brain is frequently performing active tasks, even during “rest”. A few studies have suggested personally relevant planning as one such task, which involves activity in the autobiographical memory system (Baird et al. 2011) and executive control systems (Christoff et al. 2009; Smallwood et al. 2012). Alternatively, common self-reported experiences of engaging in problem solving during rest (Smallwood and Schooler 2006) may also require increased activity in areas such as the FPN. Additionally, one might expect periodic activations in motor and sensory systems that are largely involved with passive attending to sensorimotor events (Fox et al. 2006). These “task-positive” networks are present in the State A activation pattern, suggesting State A might be important in a full explanation of mind wandering. However, future work will be required to assess the specific tasks that are performed during unconstrained resting-state mind wandering. Also of note, State B was present for a substantial portion of time during task performance. These results suggest there is a balance between these two states – with only moderate shifting from this homeostatic baseline – regardless of outward behavioral state.

### Limitations

The present study involves several limitations that will be important for future studies to address. For instance, with fMRI the choice of a representative “baseline” for activation analyses is a complex issue. For standard task analyses, this is often circumvented using inter-task rest periods as the baseline to compare across task conditions. This choice has several issues, especially given that some regions are more active during rest than task (Stark and Squire 2001). Yet even this imperfect baseline choice is unavailable when investigating activations during resting states. Building on common practice in the fMRI FC literature, we used each region’s (or voxel’s) time series mean as baseline. This is equivalent to using the spatial average across all whole-brain images in an fMRI run as the baseline, which removes brain anatomy and other potentially spurious similarities between brain images at individual time points. However, differences in the choice of baseline may influence the amplitude of the activations we observe, which in turn may affect the interpretability of activation patterns. Notably, this would be a problem for methods with a true baseline as well (e.g., positron emission tomography), given that without temporal de-meaning the spatial activation patterns could mostly reflect anatomy (and so would always be highly similar to each other). However, the majority of our analyses and findings are rooted in correlation-based methods, which are designed to remove the influence of baseline (or linear scaling) shifts in signals. Thus, our results may be less influenced by baseline issues than other alternative approaches. While the choice of a valid baseline remains an important problem to explore in fMRI research, its effect is likely minimized by the analytical approaches used in this study.

The dimensionality of the feature space, downscaling from approximately 60,000 grey-matter voxels to 264 brain regions, is another potential limiting factor for this study. While signal-to-noise is likely increased with a reduction in dimensionality (through averaging related signals within brain regions), meaningful information at the voxel level can also be lost. We focused on the scale of brain regions to take advantage of higher signal-to-noise, and also for computational tractability. Importantly, we found that the conclusions drawn at the region level generalized to the voxel level, based on mapping the time points assigned to each brain state to the voxel level (Figure 1C). It will be important, however, for future studies to perform our dMVPA approach (or related methods) directly on voxel patterns to assess potentially more detailed (but potentially noisier) activation pattern brain states.

### Conclusion

We used a novel dMVPA approach combining insights from multivariate activation methods as well as graph theoretical methods to identify and characterize activation pattern brain states during resting-state and task-state fMRI. This provided three primary benefits. First, relative to related M/EEG dMVPA approaches, the increased spatial accuracy of fMRI facilitated identification of activation patterns. Second, use of both resting-state and task-state data across 98 subjects allowed for an especially comprehensive identification of human brain activation states. Third, a focus on fMRI activation amplitude patterns allowed us to begin to interpret even spontaneous activations in terms of specific functions, given the vast task fMRI activation literature and associated cognitive functions. Together these benefits revealed a hierarchical organization of brain states with shared features across rest periods and task performance. It will be important for future studies to build on these results, improving our understanding of the many brain states that are entered spontaneously and as a result of social interactions (e.g., cognitive task instructions) along with their relation to ongoing brain network dynamics.

## ACKNOWLEDGEMENTS

We would like to thank Patrick Shafto, Bharat Biswal, and Bart Krekelberg for helpful conversations during preparation of this manuscript.

